# Pathogenic tau induces an adaptive elevation in mRNA translation rate at early stages of disease

**DOI:** 10.1101/2024.02.19.581061

**Authors:** Gabrielle Zuniga, Sakie Katsumura, Jasmine De Mange, Paulino Ramirez, Farzaneh Atrian, Masahiro Morita, Bess Frost

## Abstract

Alterations in the rate and accuracy of messenger RNA (mRNA) translation are associated with aging and several neurodegenerative disorders, including Alzheimer’s disease and related tauopathies. We previously reported that error-containing RNA that are normally cleared via nonsense-mediated mRNA decay (NMD), a key RNA surveillance mechanism, are translated in the adult brain of a *Drosophila* model of tauopathy. In the current study, we find that newly-synthesized peptides and translation machinery accumulate within nuclear envelope invaginations that occur as a consequence of tau pathology, and that the rate of mRNA translation is globally elevated in early stages of disease in adult brains of *Drosophila* models of tauopathy. Polysome profiling from adult heads of tau transgenic *Drosophila* reveals the preferential translation of specific mRNA that have been previously linked to neurodegeneration. Unexpectedly, we find that panneuronal elevation of NMD further elevates the global translation rate in tau transgenic *Drosophila*, as does treatment with rapamycin. As NMD activation and rapamycin both suppress tau-induced neurodegeneration, their shared effect on translation suggests that elevated rates of mRNA translation are an early adaptive mechanism to limit neurodegeneration. Our work provides compelling evidence that tau-induced deficits in NMD reshape the tau translatome by increasing translation of RNA that are normally repressed in healthy cells.

## INTRODUCTION

The tau protein is best known for its canonical role as a microtubule stabilizer within axons of healthy neurons^1^. Deposition of insoluble tau species is a neuropathological hallmark of over 20 neurodegenerative disorders^2–4^, collectively termed “tauopathies,” including Alzheimer’s disease. In brains of patients with Alzheimer’s disease and models of tauopathy, tau is subject to an array of posttranslational modifications that prevent binding of tau to microtubules and promote tau co-aggregation with methylated RNA^5^, RNA-binding proteins^6–8^, splicing components^9,10^, and translation machinery^11,12. 2–4^ New research points to the involvement of RNA processing defects in tauopathy, including aberrant RNA quality control and translational control of mRNA^13–17^. Advances in proteomic methodologies, including the development of polysome and ribosome profiling, make it possible to interrogate translation in detail to identify novel, alternatively processed protein products in tauopathy that may represent new potential therapeutic targets for intervention.

Changes in whole-organism translational activity are associated with aging, the greatest risk factor for the development of Alzheimer’s disease. Global translation peaks at young ages then steadily declines over time^18–22^. Neurons have higher rates of translation compared to other cell types, increasing their vulnerability to shifts in overall translation patterns^23^. Neurons also require a readily releasable pool of ribosomes to continuously remodel their proteome in response to external stimuli or experiences that strengthen or form new synaptic connections and store information as memories^24^. Recent studies report that ribosomal protein expression is reduced in late-stage human Alzheimer’s disease brain tissue^15,16,25,26^, and that tau causes a reduction of global translation in mouse models of tauopathy^14,16^. While these studies support a model in which age-associated changes in mRNA translation and protein synthesis drive memory loss in neurodegenerative diseases such as Alzheimer’s disease^27^, we currently lack mechanistic detail connecting changes in mRNA translation to tau-induced neurodegeneration.

NMD is a highly conserved RNA surveillance mechanism that safeguards the transcriptome by clearing aberrant, error-containing RNA as well as many “normal,” physiological transcripts^28^. Neurons have heightened alternative splicing compared to other cell types, providing a means of increasing intronic RNAs in the neuronal transcriptome^29,30^ that undergo a high level of turnover via NMD. Intron retention is highly prevalent in brain tissue from older patients, patients with Alzheimer’s disease^29^, and in a *Drosophila* model of tauopathy^9,13^, suggesting that the aging and degenerating brain is enriched for intron-containing transcripts that, if not cleared by NMD, could be translated into aberrant protein products that are potentially toxic to the cell. In line with these data, we recently reported that NMD is at a deficit in brains of *Drosophila* expressing the *R406W* mutant form of human tau (tau^R406W^)^13^, which is associated with autosomal dominant frontotemporal dementia^31^,and that normal and error-containing RNA accumulate within lobulations and tunnel-like invaginations of the nuclear envelope^2^. Nuclear invagination and blebbing also occur in human brain affected by Alzheimer’s disease or primary tauopathy^32,33^, iPSC-derived neuron models of tauopathy^34^, as a consequence of induced expression of tau in HEK293^8^ and neuroblastoma cells^35^, and upon induced tau multimerization in primary neurons^36^. Using a transgenic reporter of NMD activity, we found that RNA harboring an intron in the 3’ UTR escape NMD, accumulate within nuclear invaginations and adjacent to nuclear blebs, and become translated into protein in tau^R406W^ transgenic *Drosophila*^13^.

In the current study, we find that newly-synthesized peptides and ribosome-like structures are localized within nuclear envelope invaginations in tau^R406W^ transgenic *Drosophila*, and that overexpression of either tau^R406W^ or tau^WT^ (which models sporadic tauopathies) significantly enhances relative rates of global mRNA translation in *Drosophila* prior to overt neurodegeneration. Using polysome profiling in parallel with RNA sequencing, we identify a subset of mRNA transcripts that have been previously implicated in neurodegenerative disorders^37–41^ that appear to be most affected by elevated rates of global translation in tau^R406W^ transgenic *Drosophila*. Our genetic and biochemical analyses reveal that panneuronal RNAi-mediated reduction of NMD core factor Upf1 enhances global translation rate in the absence of tau, suggesting that failure to clear error-containing RNA can lead to elevated translation. Unexpectedly, we find that increasing NMD via panneuronal overexpression of NMD core factor Upf1 further elevates the translation rate in tau transgenic *Drosophila*, as does treatment of flies with rapamycin, an inhibitor of mammalian target of rapamycin complex 1 (mTORC1). As Upf1 overexpression and rapamycin have previously been reported to suppress tau neurotoxicity in tau^R406W^ transgenic *Drosophila*^13,42–44^, these data point toward a neuroprotective effect of tau-induced elevation of translation early in disease. Overall, our studies uncover a previously unrecognized relationship between tau, nuclear pleomorphism, NMD and translation relevant to the early prodromal stages of neurodegenerative tauopathies, when therapies with disease-modifying potential are most likely to have the strongest clinical impact.

## METHODS

### Drosophila genetics

*Drosophila* were fed a standard diet (Bloomington formulation: 1.5% yeast, 6.6% light corn syrup, 0.9% soy flour, 6.7% yellow cornmeal, 0.5% agar) at 25°C in a 12 hour light/dark cycle for all aging and crossing experiments. The *elav*-*GAL4* driver fly line (Bloomington Drosophila Stock Center) was used to panneuronally express transgenes or small hairpins for RNAi-mediated knockdown of transcripts in *Drosophila*. *UAS-SCA3* transgenic *Drosophila* were provided by Dr. Nancy Bonini^47^. Previously described^13,45,46^ *UAS-tau^R406W^* and *UAS-tau^WT^* transgenic *Drosophila* were provided by Dr. Mel Feany^46^. Full genotypes, insertion chromosome, stock number and sources are listed in **Supplemental Table 1**.

### Immunofluorescence

*Drosophila* brains were dissected in ice-cold PBS and fixed in 4% paraformaldehyde for 30 minutes followed by blocking with PBS plus 2% milk and 0.2% Triton (PBSTr) for at least 30 minutes. Brains were incubated with primary antibodies detecting puromycin and lamin in PBSTr+2% milk overnight at room temperature. Secondary detection was performed using Alexa Fluor-conjugated secondary antibodies, incubated with the sample for two hours before mounting slides with Fluoromount G with DAPI (Southern Biotech). All images were processed as a single slice using a Zeiss confocal microscope and analyzed using ImageJ software. Primary antibodies, sources and concentrations are included in **Supplemental Table 2**.

### Electron microscopy

Fly heads were fixed in phosphate buffered 4% formaldehyde and 1% glutaraldehyde for at least two hours followed by osmium tetroxide treatment for 30 minutes. The samples were then dehydrated through a graded series of ethanol and sent to the Electron Microscopy Laboratory in the Department of Pharmacology and Laboratory Medicine at the University of Texas Health San Antonio for further processing. Each section was embedded with resin in a flat mold for correct orientation and then polymerized in an 85°C oven overnight. Thin sections were captured on copper grids using a JEOL 1400 Transmission Electron Microscope.

### Drug treatment

Puromycin (BP2956, Fisher Scientific) was diluted in water to a final concentration of 90 mM and added to standard fly food at a final concentration of 10 mM. Water was used as a vehicle control. *Drosophila* were collected after eclosion and transferred to food containing puromycin 24 hours prior to being frozen or fixed.

### Immunoblotting

Frozen *Drosophila* heads were homogenized in 15 µL RIPA buffer containing both protease and phosphatase inhibitors (ThermoScientific) and total protein content was measured using a BCA protein assay kit (ThermoScientific). All samples were diluted to the same protein concentration using 2X Laemmli buffer (Sigma-Aldrich), boiled for 10 minutes, and briefly centrifuged. Equal amounts of protein (12-15 µg) were run on 10% SDS-PAGE precast gels. Protein samples were transferred to nitrocellulose membranes, checked for equal loading using Ponceau S staining, and blocked in 2% milk in PBS with 0.05% Tween for 10-30 minutes. Membranes were immunoblotted overnight at 4°C with primary antibody. An antibody against β-actin was used to assess equal protein loading. After washing, membranes were incubated with HRP-conjugated secondary antibodies and developed with an enhanced chemiluminescent substrate (ThermoFisher). For quantitative analyses, densitometry was performed and the relative level of each protein relative to actin was calculated using ImageJ software.

### Polysome profiling

Polysome profiling analysis was carried out as previously described, with a few exceptions^47^. Approximately 300-500 heads from control and tau^R406W^ transgenic *Drosophila* at day 10 of adulthood were frozen on dry ice, separated using a sieve, and homogenized in lysis buffer (10 mM Hepes-KOH, 150 mM KCl, 5 mM MgCl2, 0.5 mM DTT, RNase inhibitors, 100 µg/ml cycloheximide, Protease inhibitors). 0.5% NP40 was added to the lysate followed by centrifugation at 2000 x *g* for ten minutes at 4°C. After five minutes of incubation on ice, the samples were centrifuged at 20,000 x *g* for ten minutes at 4°C. After 20 minutes of incubation on ice,the supernatant was transferred and OD_260_ was used to quantify RNA content using a Nanodrop 8000 spectrophotometer (Thermo Scientific). An equal amount of RNA based on OD_260_ was added to each gradient. Sucrose gradients were prepared beforehand, stored at −80°C, and thawed at 4°C at least two hours prior to polysome isolation. Sucrose gradients containing lysate were run at 35,000 rpm for two hours at 4°C using the SW40Ti rotor in a Beckman Coulter Optima L-80 ultracentrifuge. For RNA analysis, the chasing solution (60% [w/v] sucrose bromophenol blue) was run through the gradient with the pump set at 1.5 ml/min and fractions were collected every 30 seconds (Brandel). A_254_ was measured for each fraction.

The libraries were prepared and sequenced by the Genome Sequencing Facility at Greehey Children’s Cancer Research Institute at the University of Texas Health San Antonio. RNA sequencing was carried out using the Illumina NovaSeq 6000 system with an average of 30 million 100 base pair paired-end reads per sample. Fastp (v0.23.2) was utilized for trimming and quality control of reads. Trimmed reads were aligned to the *Drosophila melanogaster* Flybase v6.43 genome using STAR (2.7.10a) with default parameters. Read counts were obtained with STAR. Changes in translation efficiency between control and tau^R406W^ *Drosophila* were identified using the anota2seq R package^50^ using rlog normalized counts and filtering out genes with no coverage. Genes were classified as having a change in translation efficiency if they passed the threshold suggested by anota2seq maximum with an adjusted p-value of 0.15 and a slope p-value of 0.1. For anota2seq, using an adjusted p-value of 0.15 to identify mRNA in each regulatory group (translational efficiency, translational buffering, and translational equivalency) decreases the number of false negatives compared to an adjusted p-value of 0.05^48^.

### Statistical analyses

All experiments utilized an equal number of male and female *Drosophila.* When possible, samples were randomized and investigators were blinded to genotype/treatment. Biological replicates, reported as n, were based on previous publications^40,51^. Statistical analyses were performed using unpaired Student’s t-test when comparing two samples. ANOVA with post-hoc test was used when comparing multiple samples.

## RESULTS

### Active translation occurs within tau^R406W^-induced nuclear invaginations

We took advantage of the short life cycle, low maintenance cost, and powerful genetics available to the *Drosophila* model system to investigate the relationship between nuclear pleomorphism and protein synthesis in *in vivo* models of tauopathy. Neuron-specific expression of human tau^R406W^ or wildtype human tau (tau^WT^) in *Drosophila* drive neurodegeneration through mechanisms including disruption of the lamin nucleoskeleton^32^, widespread decondensation of constitutive heterochromatin^45^, intron retention^9^, DNA damage^49^ and oxidative stress^50^, to name a few, that are conserved in human Alzheimer’s disease. As tau^R406W^ transgenic *Drosophila* provide a moderate level of toxicity at day ten of adulthood that is well-suited for genetic analyses.

To resolve the subcellular localization of ribosomes, the active players in protein translation, we imaged brain of ten-day-old tau^R406W^ transgenic *Drosophila* using electron microscopy (EM). We observe a striking accumulation of individual electron-dense beads and string of beads that resemble ribosomes and mRNA-bound polysomes, respectively, adjacent to nuclear blebs and within involutions of the nuclear envelope (**Fig. 1A**). These results suggest that the cytoplasmic core of nuclear invaginations have the capacity for local protein synthesis.

**Figure 1.**
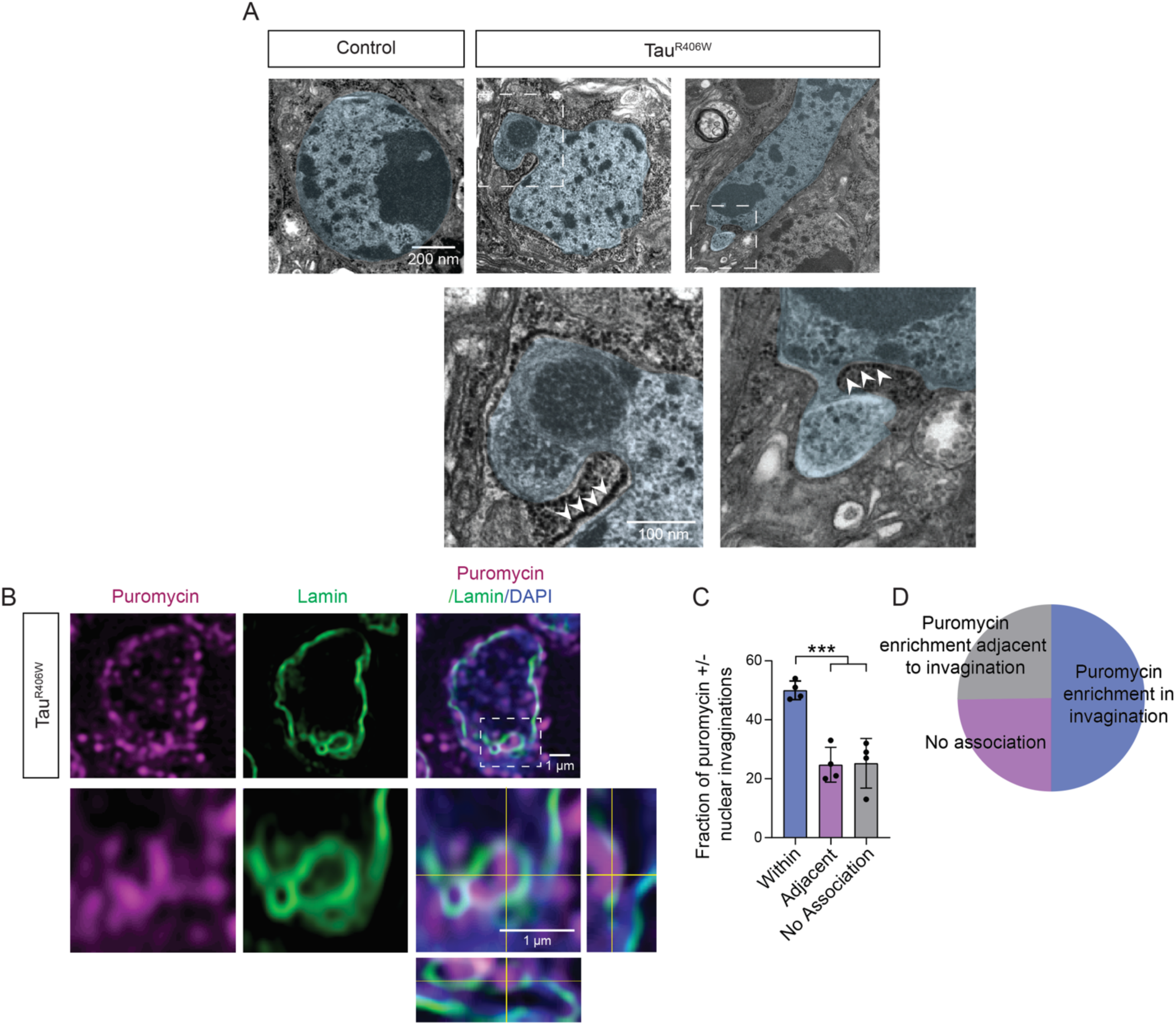
Ribosomes and translational products are localized within nuclear envelope invaginations in brain of tau^R406W^ transgenic *Drosophila*. A) Representative images show accumulation of electron-dense particles within nuclear envelope invaginations in brain of tau^R406W^ transgenic *Drosophila* as visualized via EM. Nuclei and nuclear blebs are artificially colored blue for visualization. **B)** Immunofluorescence-based detection of puromycylated peptides suggest that active translation occurs within nuclear invaginations. Orthogonal view of the zoom image depicted shows bright puromycin-positive foci clearly within invaginations, outlined by the lamin nucleoskeleton. **C)** Quantification of the percentage of puromycylated peptides within, adjacent, or not associated with nuclear invaginations, as shown in pie chart **(D)**. n=4 biological replicates. p***<0.001, one-way ANOVA. Error bars=SEM. All assays were performed at day 10 of adulthood.

Having localized electron-dense particles resembling ribosomes within nuclear invaginations of tau^R406W^ transgenic *Drosophila*, we next asked if RNA is actively translated into protein within nuclear invaginations. To detect translating ribosomes with single-cell resolution, we utilized puromycin, an antibiotic and tyrosyl-tRNA analog that readily incorporates into newly synthesized proteins by occupying the acceptor site of ribosomes^51^. We fed tau^R406W^ transgenic *Drosophila* puromycin for 24 hours and visualized the relationship between translating ribosomes and nuclear invaginations and blebs by co-labeling *Drosophila* brains with antibodies detecting puromycin and lamin. In tau^R406W^ transgenic *Drosophila*, we detect focal accumulation of puromycylated peptides (**Fig. 1B**) within approximately half of nuclei harboring invaginations (**Fig. 1C, D**). Overall, these data provide compelling evidence that ribosomes actively engage in protein synthesis within tau-induced nuclear invaginations.

### Tau^R406W^-induced increase in active translation precedes significant neurodegeneration and lifespan decline

Given that the level of puromycylated peptides can serve as a proxy for measuring relative rates of global translation^52^, we adapted the puromycin-based nonradioactive surface sensing of translation (SUnSET) assay^52^ to quantify mRNA translation rate in *Drosophila* heads at day ten of adulthood. Previously used for quantifying translation rate in cultured cells, the SUnSET assay relies on the ability of puromycin to incorporate into elongating polypeptide chains at low concentrations without affecting overall translation and inducing cell death^52^. To determine whether overall translation rate is affected by tau, we measured levels of puromycin using a fluorescein-labelled anti-puromycin antibody in brain of puromycin-fed control and tau^R406W^ transgenic *Drosophila*. We find a robust increase in puromycin signal in the cortex of tau^R406W^ transgenic *Drosophila* brain at day ten of adulthood (**Fig. 2A, B**), suggesting that panneuronal expression of pathogenic tau enhances the rate of global translation.

**Figure 2.**
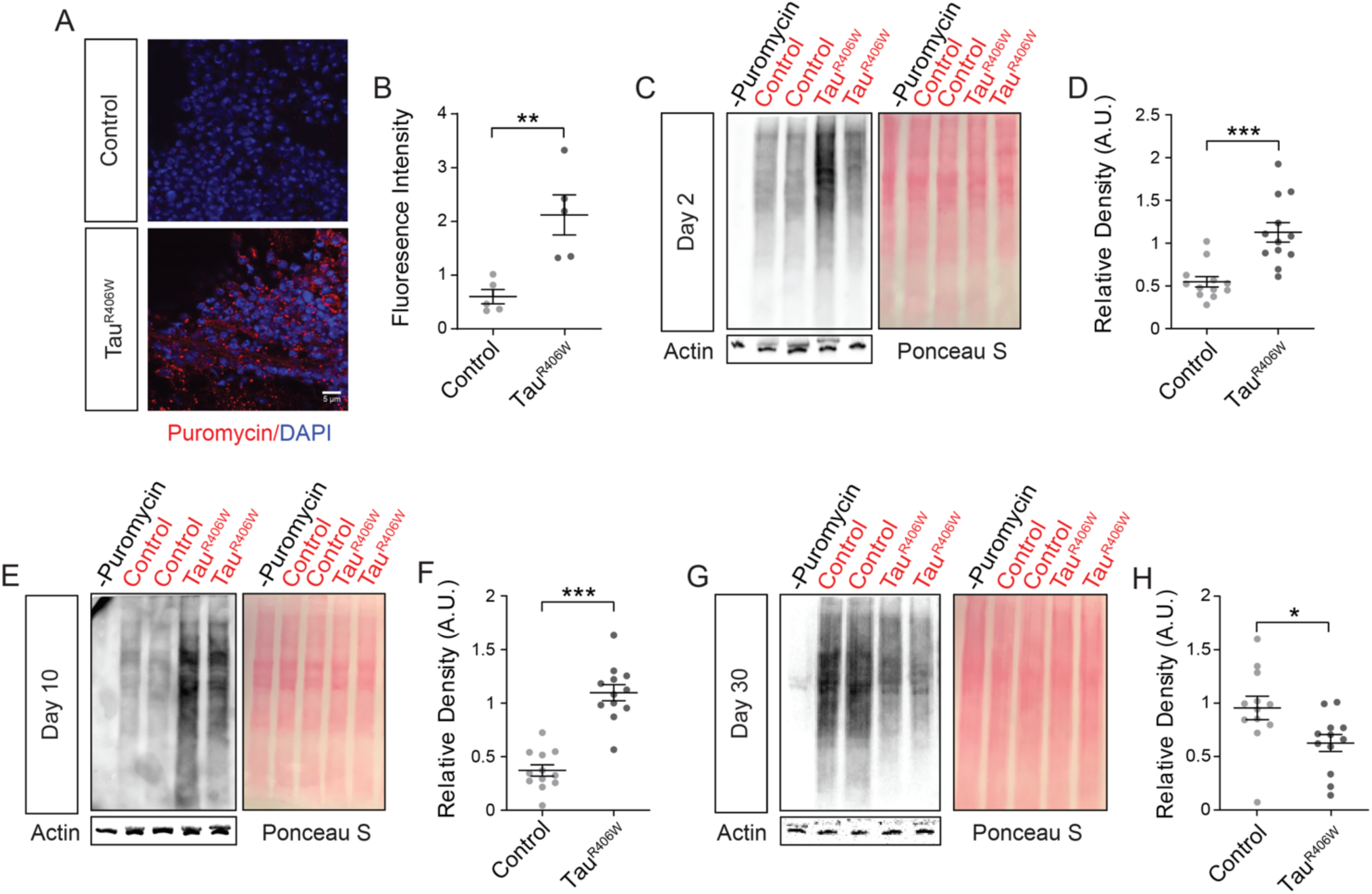
Global translation rate is increased prior to significant neurodegeneration in tau^R406W^ transgenic *Drosophila*. **A)** Representative image of puromycylated peptides in the cortex of control and tau^R406W^ transgenic *Drosophila*. **B)** Immunofluorescence-based analyses of puromycin within the *Drosophila* brain reveal significantly increased puromycin levels in tau^R406W^ transgenic *Drosophila* compared to controls. n=4 biological replicates per genotype**. C-F)** Western blot analysis of puromycin levels relative to Ponceau S staining reveal that panneuronal expression of transgenic human tau significantly elevates global translation rate in heads of tau^R406W^ transgenic *Drosophila* at day two **(C, D)** and ten **(E, F)** of adulthood, but significantly depletes global translation rate at day 30 (**G, H**). ^55^n=12 biological replicates per genotype, per age. *p<0.05, **p<0.01, ***p<0.001, unpaired t-test. Error bars=SEM.

As a complementary approach, we next used our modified SUnSET assay to quantify translation rate in *Drosophila* by western blotting. After treatment with puromycin, puromycin-labeled nascent proteins appear as a smear upon incubation of western blotting membranes with an anti-puromycin antibody. Based on previous work showing that inactivation of the eukaryotic translation initiation factor 4E binding protein (eIF4EBP) is sufficient to activate translation *in vivo*^53–55^, we first quantified the rate of global translation in *Drosophila* with panneuronal RNAi-mediated reduction of *Thor*, the *Drosophila* ortholog of human eIF4EBP 1 and 2, as a positive control for elevated translation. We find that *Thor* knockdown increased levels of puromycin relative to puromycin-fed controls and background (no puromycin treatment) (**Supplemental Fig. 1**), establishing the fidelity of our protocol.

Turning our focus to tauopathy, we find that tau^R406W^ transgenic *Drosophila* have increased global translation rate relative to control at day two of adulthood (**Fig. 2C, D**), which is prior to significant neurodegeneration^46^. Elevated translation is sustained in tau transgenic *Drosophila* at day ten of adulthood (**Fig. 2E, F**), a point at which neurodegeneration can be detected^46^ but survivorship is not significantly reduced^56^ in this model. We find that global translation rate at day 30 of adulthood, which nears the maximum lifespan of this model, is significantly reduced in heads of tau^R406W^ transgenic *Drosophila* compared to controls (**Fig. 2G, H**), which is consistent with reduced ribosomal protein expression in late-stage human Alzheimer’s disease brain tissue^14,57^. Our findings suggest that expression of pathogenic tau drives increased global translation early in the disease process while later bringing about a significant decline in global translation below that of age-matched controls, thus highlighting the importance of analyzing cellular phenotypes across disease stages.

### Increased translation is not a general feature of neurodegeneration in Drosophila

Having found a significant increase in mRNA translation rate at early disease stages in tau^R406W^ transgenic *Drosophila*, we next asked if tau-induced elevation of mRNA translation in the *Drosophila* brain is a specific consequence of the *R406W* mutation. Tau^WT^ aggregates in brains of patients with sporadic tauopathies, including Alzheimer’s disease^58^, and drives neurodegeneration in fly and mouse models of tauopathy, although less aggressively than mutant forms of tau^46^. At ten days of adulthood, we find that the rate of global translation is significantly increased in tau^WT^ transgenic *Drosophila* (**Fig. 3A, B**), indicating that increased translation rate is conserved in a *Drosophila* model of sporadic tauopathy, and is not a specific feature of the *R406W* mutation.

**Figure 3.**
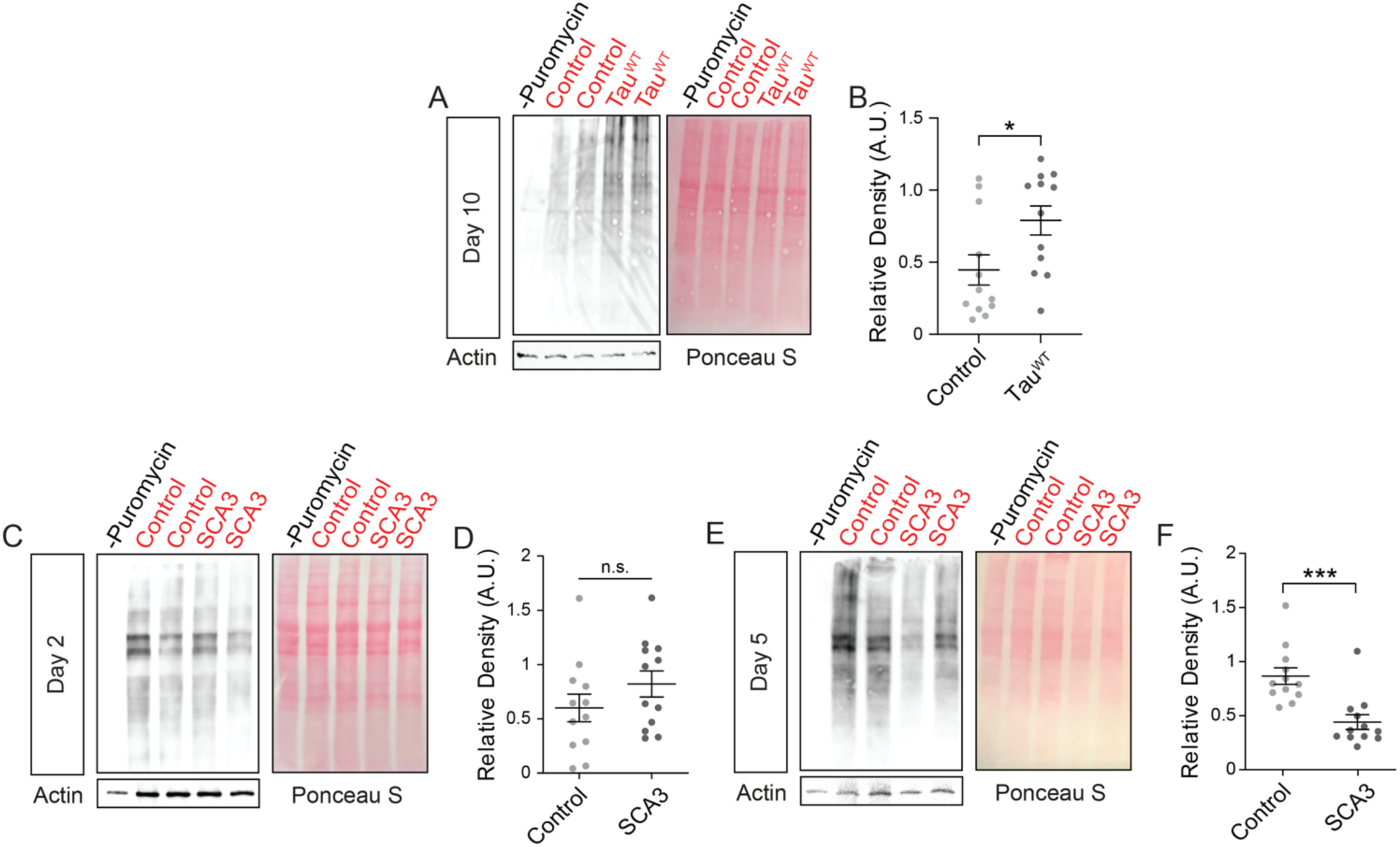
Increased mRNA translation is a general feature of tauopathies. **A)** Panneuronal expression of tau^WT^ significantly increases global translation rates based on western blot. **B)** Quantification was determined relative to Ponceau S. **C-F)** Global translation rate in heads of SCA3 transgenic *Drosophila* is unchanged at day two **(C, D)**, but is significantly reduced at day five **(E, F)** compared to age-matched controls. n=12 biological replicates per genotype. *p<0.05, **p<0.01, unpaired t-test. Error bars=SEM.

To determine if increased translation rate is a common feature of neurodegeneration in *Drosophila*, we quantified the global translation rate in a *Drosophila* model of spinocerebellar ataxia type 3 (SCA3), also known as “Machado Joseph Disease” (MJD), which panneuronally expresses a truncated form of the human *MJD1* gene with an expanded repeat of 78 glutamines (MJDtr-78)^59^. We analyzed translation rate at days two and five of adulthood, as neurodegeneration in this model is relatively more aggressive than tau^R406W^ transgenic *Drosophila.* While translation rates are unchanged between SCA3 and control *Drosophila* at day two of adulthood (**Fig. 3C, D**), translation rates are significantly reduced in the SCA3 model by day five of adulthood (**Fig. 3E, F**). Together, these data suggest that elevated translation rate at earlier disease stages is not a general feature of neurodegeneration, but that reduced translation rate at later stages of disease may be a general consequence of neurodegeneration.

### Polysome profiling identifies differentially translated mRNAs in tau^R406W^ transgenic Drosophila

As our experiments based on puromycin labeling suggest that global translation rate is increased at early and mid-stages of disease in tau^R406W^ transgenic *Drosophila,* we next sought to identify transcripts that are actively translated. We combined RNA sequencing and polysome profiling on the same sample in order to 1) identify actively-translated transcripts and 2) determine translation efficiency of individual transcripts in heads of 10-day-old tau^R406W^ transgenic *Drosophila* compared to controls.

After isolating ribosomal fractions from control and tau^R406W^ transgenic *Drosophila* heads (**Fig. 4A**), we determined the translational status of all mRNA by comparing the proportion of efficiently translated mRNA (defined as RNA associated with three or more ribosomes (polysomes)) to total RNA (raw counts are provided in **Supplemental Table 3**). Translation efficiency is calculated as the number of ribosomes associated with an mRNA (average ribosome footprint density) divided by the total abundance of that mRNA. Polysome profiling analysis requires parallel mRNA sequencing in order to discriminate between three scenarios: 1) Polysome-associated mRNA exceeds total mRNA (high translational efficiency) or is less than total mRNA (low translational efficiency) (**Fig. 4B**, left panel), 2) Polysome-associated mRNA is unchanged despite altered mRNA abundance (translational “buffering”) (**Fig. 4B**, middle panel), and 3) Polysome-associated mRNA is equivalent to total RNA levels, such that changes in translation match corresponding alterations in mRNA levels (translational equivalency) (**Fig. 4B**, right panel). As previously described^60^, heads from hundreds of flies are required for polysome profiling, making the procedure likely to identify only the largest changes in translational efficiency.

**Figure 4.**
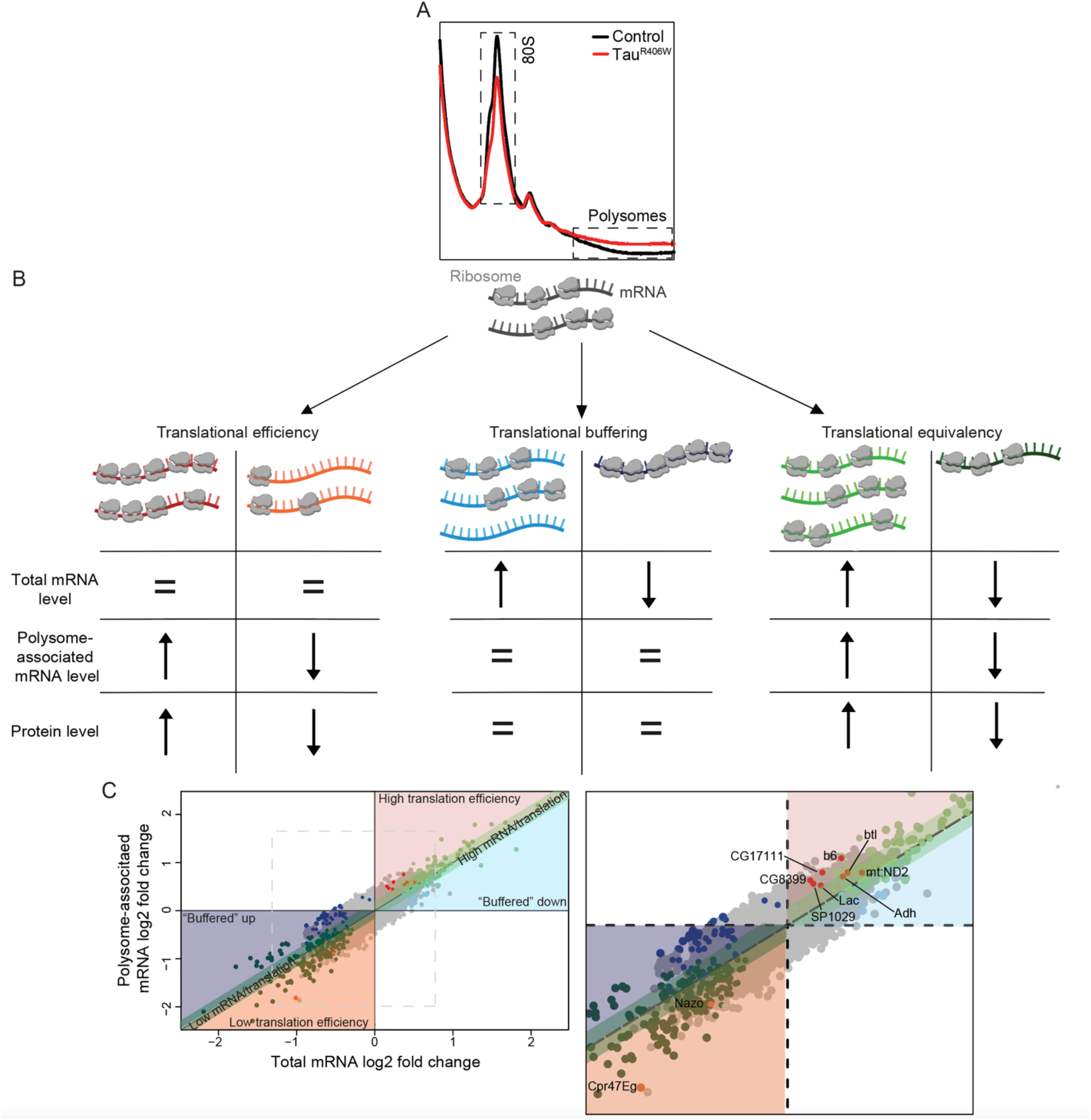
Polysome profiling reveals a subset of efficiently translated mRNA in tau^R406W^ transgenic *Drosophila* heads. **A)** Polysome profiles obtained from heads of control (black line) and tau^R406W^ transgenic *Drosophila* (red line) at day 10 of adulthood. n=5 biological replicates per genotype. **B)** Schematic representation of gene expression regulation that occurs at the level of translation. Left panel: changes in polysome-associated mRNA levels affecting the proportion of total mRNA being translated, representing changes in “translation efficiency”; Middle panel: “translational buffering” in which unaltered polysome-associated mRNA levels offset changes in mRNA levels; Right panel: “translational equivalency” in which total mRNA abundance is equivalent to polysome-associated mRNA levels. Colors represent key to shading and dots in scatterplot below. **C)** Scatterplot of RNAs with differential “translational efficiency” in tau^R406W^ transgenic *Drosophila* compared to controls (significantly high translation efficiency, red dots; significantly low translation efficiency, orange dots), “translational buffering” reflecting unaltered mRNA polysome association despite changes in mRNA levels (high mRNA levels, light blue dots; low mRNA levels, dark blue dots), and “translational equivalency” reflecting similar levels of total mRNA and polysome-associated mRNA (high total/polysome-associated mRNA, light green dots; low total/polysome-associated mRNA levels, dark green dots). Zoom image depicted with differentially translated mRNA labeled. n=4 biological replicates per genotype. Adjusted p-value<0.15.

When comparing tau^R406W^ transgenic *Drosophila* to control, we identify ten mRNA that fit the first scenario (**Fig. 4C**), which indicates an alteration in translational efficiency. Among genes with higher than expected polysome association, i.e. efficiently translated RNAs, several have been previously associated with Alzheimer’s disease^37,38^, tau pathology^39–41,61^ and various diseases of the brain^62–66;^ genes that when knocked down or overexpressed in a mutant form can drive neurodegeneration in animal models^67,68^; a gene that genetically modifies tau-induced neurodegeneration in *Drosophila*^69^; and genes that are predicted to be protective against neurodegeneration^69,70^ (**Supplemental Table 4**). Proteins encoded by *ferric chelate reductase 1 (FRRS1)*, *15-hydroxyprostaglandin dehydrogenase (HGPD-15)*, and *neuronal pentraxin 2 (NPTX2)*, the human orthologs of *ascorbate ferrireductase*, *adh*, and *b6*, respectively, are increased in brain of mouse models of Alzheimer’s disease^71^ and human patients with Alzheimer’s disease^37,40^ (**Supplemental Table 4**), suggesting that the translational status of mRNA in heads of tau^R406W^ transgenic *Drosophila* is relevant to the proteome in vertebrate and human tauopathy. Taken together, these findings suggest that elevated levels of proteins previously identified and implicated in various models of Alzheimer’s disease and related tauopathies may be a consequence of preferential mRNA translation.

### Reduced clearance of RNA by NMD drives increased translation

We next took steps to determine the mechanism by which tau drives an increased global translation rate. Previous work in the *Drosophila* tau^R406W^ and tau^WT^ models suggest that pathogenic tau causes a toxic deficit in NMD, resulting in accumulation of NMD-target transcripts that are actively translated into protein in the cytoplasm and within nuclear envelope invaginations and blebs^13^. The classic model posits that NMD of a particular mRNA takes place during translation and is activated when up-frameshift factors (Upfs) accumulate on the 3’ UTR. Suppressor of <u>m</u>orphological defects on <u>g</u>enitalia 1 (smg1) and Smg5-9 are recruited to the Upf1-bound mRNA and promote phosphorylation of Upf1 and rapid degradation of the target mRNA^72^. We first hypothesized that reduced clearance of RNA by NMD increases translation in *Drosophila*. To test this hypothesis, we reduced expression of NMD core factor Upf1 in neurons of *Drosophila* via RNAi and quantified translation rate using our modified SUnSET assay. We find that panneuronal knockdown of Upf1 increases the rate of global translation in *Drosophila* heads (**Fig. 5A, B**), suggesting that deficits in RNA clearance are sufficient to increase translation.

**Figure 5.**
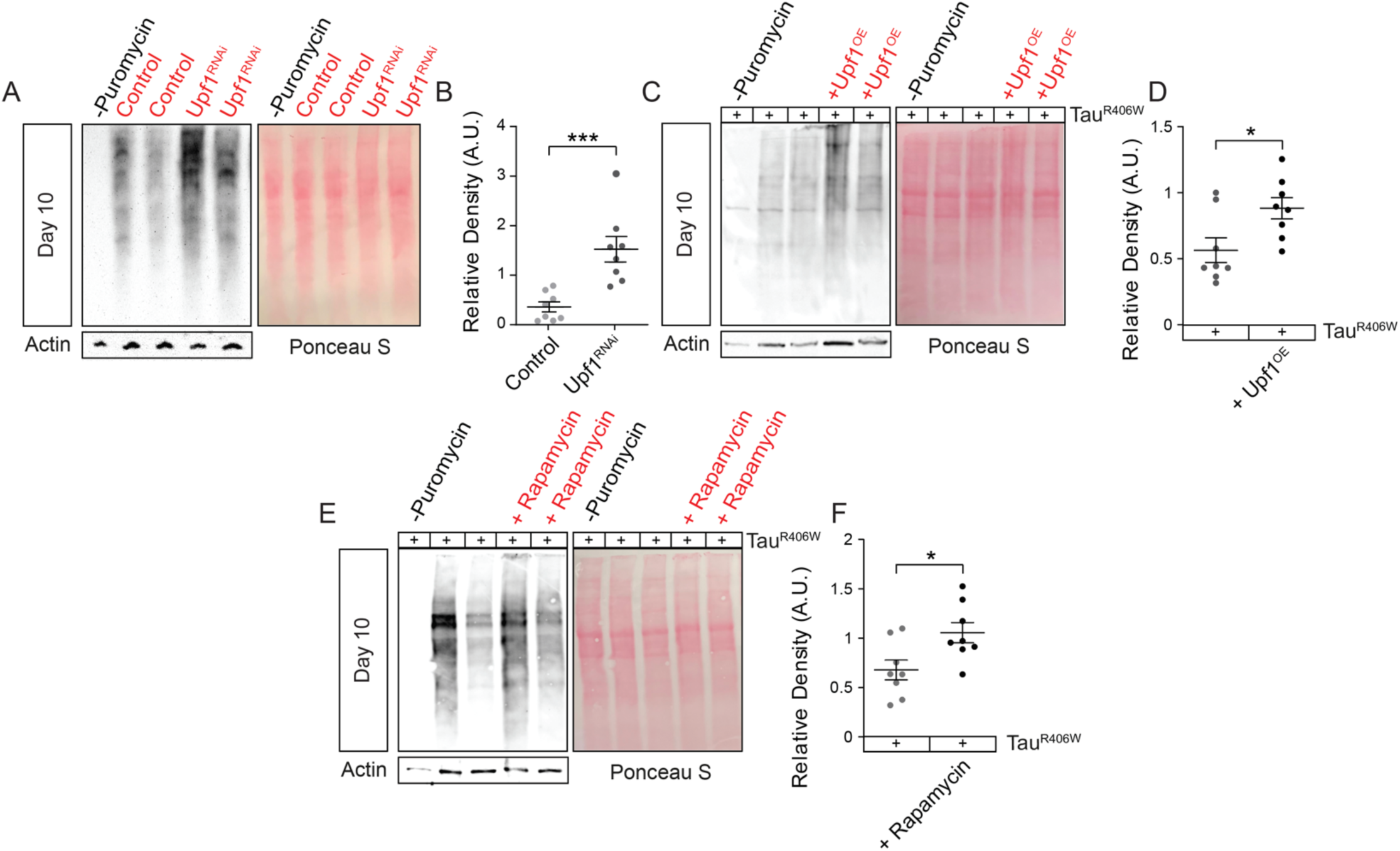
Genetic manipulation of NMD modifies global translation rates in a context-dependent manner. **A)** Western blotting reveals that panneuronal RNAi-mediated reduction of Upf1 activates global translation in *Drosophila* based on detection with puromycin **(B)**, but panneuronal overexpression of transgenic human tau and wild-type Upf1 enhances tau-induced increase in mRNA translation rates in *Drosophila* **(C, D)**. SUnSET assay reveals that chronic rapamycin treatment enhances global translation rates in heads of tau^R406W^ transgenic *Drosophila* at day ten of adulthood based on western blotting **(E)** and quantification of puromycin levels relative to Ponceau S **(F)**. n=8 biological replicates per genotype. All assays were performed at day 10 of adulthood. *p<0.05, ***p<0.001, unpaired t-test. Error bars=SEM.

Having found that reduced NMD activity drives increased rates of global translation, we expected that genetically increasing NMD in tau^R406W^ transgenic *Drosophila* would reduce translation. Life-long suppression of translational activity can significantly reduce neurodegeneration^73^ and would explain, at least partially, the neuronal health benefits provided by overexpressing wild-type Upf1 in tau^R406W^ transgenic *Drosophila*^13^. To our surprise, we find that panneuronal overexpression of wild-type Upf1 (**Fig. 5C, D**) further enhances translation rates in day-10-old tau^R406W^ transgenic *Drosophila* compared to tau expressed alone. These results were remarkably similar to treatment with rapamycin, an inhibitor of mTOR signaling that regulates protein translation and significantly reduces tau toxicity in *Drosophila*^42–44,49^. We found that ten-day exposure to 2 µM rapamycin in food significantly enhances the rate of global translation in tau^R406W^ transgenic *Drosophila* compared to vehicle (**Fig. 5E, F**). These data suggest that NMD differentially controls translation in physiological conditions versus tauopathy, and that elevated translation at an early stage of disease is neuroprotective in tauopathy.

## DISCUSSION

Translation is a complex process that responds to diverse environmental stimuli to determine the spatial, temporal, and quantitative properties of protein synthesis. Although mRNA translation has been explored in the context of Alzheimer’s disease as early as the 1980s, these results were inconsistent^74,75^, involved a small sample size, were conducted in a dish^26^, or utilized polysomes extracted from postmortem brain tissue^57^ to quantify translation. Since spatial regulatory influences on mRNA translation are lost *in vitro*, new proteomic methodologies have been developed that leverage molecular probes to tag and isolate newly-synthesized peptides and assess mRNA translation over time in living animal models. Here we perform *in vivo* analysis of mRNA translation rate in *Drosophila* models of tauopathy, and find that the rate of global translation is elevated prior to significant tau-induced neurodegeneration. We discover that tau-induced deficits in NMD affect the translatome, and that an early increase in translation is neuroprotective in tauopathy.

Based on an *in vivo* SUnSET assay, we find that the rate of mRNA translation is elevated prior to significant neurodegeneration in *Drosophila* models of tauopathy. While puromycin-based labelling can provide a snapshot of the cellular translatome, this technique only measures nascent peptides. Recent studies report that puromycylated peptides can split from ribosomes and diffuse far from their original site of translation within seconds to minutes in live *C. elegans* gonads and in a human cell line^76^. While our use of a puromycin-specific antibody does accurately label translational products, we recognize that puromyclated peptides are spatially dynamic. Presence of ribosomes within nuclear envelope invaginations nevertheless supports the conclusion that proteins are actively translated within invaginations. Puromycin and translational machinery, including ribosomes, have also been detected within nuclear invaginations associated with motor neurons derived from patients with *C9orf*72-associated amyotrophic lateral sclerosis (ALS)^77^, human mammary epithelial cells^78^, and neurons of the arcuate nucleus in rats^79^.

We find that puromycin readily incorporates into elongating polypeptides and accumulates in the *Drosophila* brain without notable toxicity. Previous detection of nascent peptides via western blotting indicated that the rate of global translation is significantly reduced in HEK293 cells overexpressing disease-associated mutant forms of human tau, primary neurons derived from rTg4510 tau transgenic mice^26^ and the K3 mouse model of tauopathy^14^. Expression of the amino-terminal projection domain, but not the carboxy-terminal or microtubule-binding repeat domain, of mutant human tau in HEK293 cells was found to be sufficient to decrease the rate of global translation^14^. Tau is sequentially cleaved in human tauopathy brain tissue, including at the amino-terminal projection domain. The tau^R406W^ and tau^WT^ constructs used in the current study lack the amino-terminal domain. Although brains harvested from living rTg4510 mice injected with puromycin showed an overall decrease in rates of global translation at seven months, no change was detected at five months^16^. We speculate that the rate of protein synthesis is increased in the brain of puromycin-treated rTg4510 mice at earlier stages of disease, prior to significant neuronal or synaptic loss^80^. Previous work in *Drosophila* report that knockout of endogenous tau activates translation^81^. In line with our findings, Portillo et al. (2021) report that acetylation of tau increases translation, which depletes ATP stores in cells and may ultimately lead to neurodegeneration^17^.

We combine polysome profiling with next-generation sequencing to identify the translational status of transcripts in tau transgenic *Drosophila*. We find that eight of the ten mRNAs that are subject to significant differences in translational efficiency in tau^R406W^ transgenic *Drosophila* compared to controls have previously been implicated in Alzheimer’s disease^37,38^, in the context of tau pathology^39–41,61^, or neurodegeneration^67,68^. As our approach does not facilitate the identification of small changes in the translation efficiency of mRNA given the large input material required for polysome profiling (∼1,000 fly heads per sample), the more efficiently translated mRNA, while only a very small fraction, likely represent the most robust changes in response to expression of pathogenic tau. These data suggest that tau-induced increase in mRNA translation alters the tau proteome at early disease stages by increasing levels of growth promoting factors, postsynaptic proteins, enzymes associated with oxidative metabolism and protein homodimerization, and proteins affecting tau proteolysis.

We recently reported that overexpressing the NMD core factor Upf1 suppresses tau-induced toxicity^13^, a key finding that has been similarly demonstrated in fly and mouse models of ALS and frontotemporal dementia (ALS/FTD)^77,82–84^. In the context of ALS/FTD, however, expression of ALS-causing mutations modifies the ability of NMD to clear RNA by suppressing global protein translation^83,84^. In contrast, we find that loss of NMD activity is sufficient to increase the rate of global translation in the absence of transgenic tau, presumably because less RNA is cleared via NMD and are instead translated into proteins. In the presence of tau, however, we find that the rate of global translation is paradoxically increased in response to genetic activation of NMD. We speculate that genetic activation of NMD suppresses tau-induced neurodegeneration by clearing error-containing RNA prior to neurodegeneration, and later upregulating translation of specific mRNA in response to neurodegeneration.

Overall, our findings provide compelling new evidence that reduced clearance of RNA by NMD activates translation and that increased translation at an early stage of disease may be critical to suppressing tau-induced neurodegeneration. Tau has been shown to modulate translation by changing the expression of ribosomal subunits and other translational machinery^17^ or by directly binding to ribosomal proteins^16^. Here we provide evidence that tau-induced changes in NMD also mediate translational rate and may contribute, at least in part, to the tau translatome across disease stages. Based on our current findings and previous studies to date, we propose a working model in which tau-induced deficits in NMD drive higher rates of global translation at early disease stages, which consumes significant cellular energy that precipitates an acute drop in translation at later stages of disease. We have previously reported that tau-induced deficits in NMD are a pharmacologically targetable, mechanistic driver of neurodegeneration^13^. Our findings provide new evidence linking tau-induced nuclear envelope pleomorphisms to NMD and the tau translatome, and informs future research investigation into how tau-mediated changes in translation across disease stages differentially impacts neurodegeneration and clinical outcomes.

## AUTHORS’ CONTRIBUTIONS

GZ and BF developed the conceptual framework of the study. Experiments were performed by GZ, SK, PR, FA and JDM. GZ analyzed data and prepared the manuscript. BF and MM contributed to data analysis and manuscript preparation.

## FUNDING

This project was supported by R01 AG057896 (BF) and T32 AG021890, T32 GM113896, TL1 TR002647, and T32 NS082145 (GZ).

## ACKNOWLEDGEMENTS

We thank the Bloomington Drosophila Stock Center (NIH P40OD018537) for providing transgenic fly strains. Transgenic human tau fly lines were provided by Dr. Mel Feany, and flies expressing UAS-SCA3 were provided by Dr. Nancy Bonini. The authors acknowledge the Electron Microscopy Laboratory in the Department of Pharmacology and Laboratory Medicine at the University of Texas Health San Antonio for processing fly samples for electron microscopy and the Texas Advanced Computing Center (TACC) at the University of Texas at Austin for providing HPC resources that have contributed to the bioinformatic analyses reported within this paper. For library preparation and sequencing services, we acknowledge the Genome Sequencing Facility at Greehey Children’s Cancer Research Institute at the University of Texas Health San Antonio, which is supported by NIH-NCI P30 CA054174, NIH S10OD030311, and CPRIT Core Facility Award (RP220662).

## AVAILABILITY OF DATA AND MATERIALS

Polysome profiling data will be uploaded to the GEO repository upon acceptance of the manuscript.

## DECLARATION OF COMPETING INTERESTS

The authors declare they have no competing interests.

## ETHICS APPROVAL AND CONSENT TO PARTICIPATE

Not applicable.

## CONSENT FOR PUBLICATION

Not applicable.

---

**Supplemental Figure 1.**
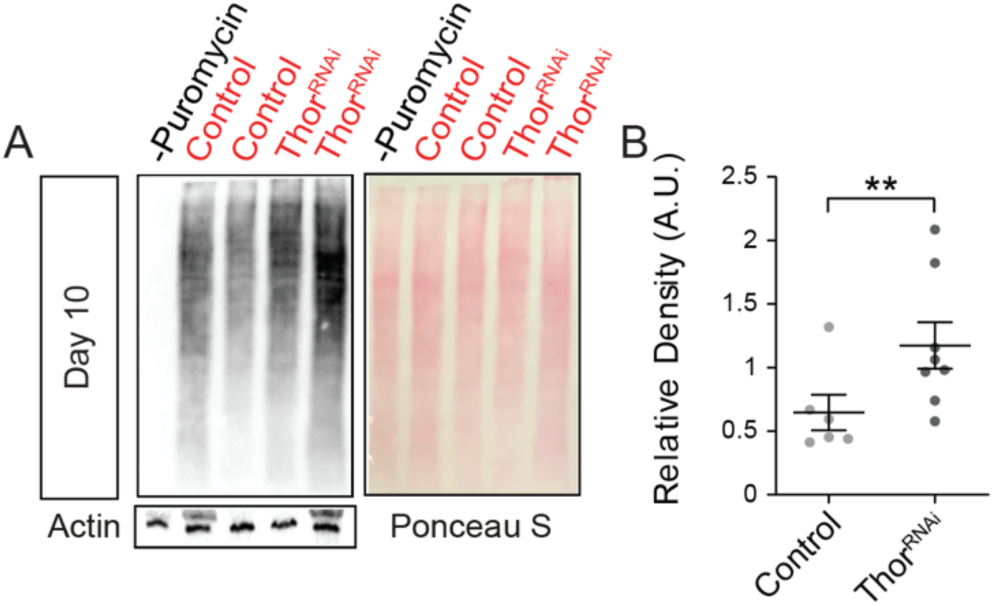
Derepression of mRNA translation significantly increases puromycin staining by western blot. **A)** Genetic reduction of Thor, *Drosophila* homologue of eIF4E binding protein, specifically in neurons of *Drosophila* fed 10 mM puromycin for 24 hours significantly increases puromycin staining by western blot relative to control, with quantification (B). p**<0.01, unpaired t-test. Error bars=SEM.

**Supplemental Table 1.** Name of *Drosophila* lines as referred to in paper as well as full genotype, insertion chromosome, stock number and source. Bloomington (Bloomington *Drosophila* Stock Center).

| Name | Full Genotype | Insertion Chromosome | Stock # | Source |
| --- | --- | --- | --- | --- |
| Upf1<sup>RNAi</sup> | y[1] v[1]; P{y[+t7.7] v[+t1.8]=TRiP.GL01485}attP2 | 3 | 43144 | Bloomington (DRSC-TRiP) |
| Upf1<sup>OE</sup> | w[*]; P{w[+mC]=UASp-GFP.Upf1}2 | 2 | 24623 | Bloomington |
| Upf2<sup>OE</sup> | w[*]; P{w[+mC]=UAS-Upf2.A}3 | 3 | 60578 | Bloomington |
| UAS-tau<sup>R406W</sup> | UAS-tau<sup>R406W</sup> | 3 |  | Mel Feany(Wittman) |
| UAS-tau<sup>WT</sup> | UAS-tau<sup>WT</sup> | 3 |  | Mel Feany(Wittman) |
| UAS-SCA3 | P{UAS-SCA3.fl-Q84.myc}7.2 | 3 |  | Nancy Bonini(Warrick) |

**Supplemental Table 2.** Antibodies used in immunofluorescence (IF) and western blotting.

| **Antibody** | **Host** | **Western** | **IF** | **Source** |
| --- | --- | --- | --- | --- |
| cTau | Mouse | 1:100,000 |  | Dako # A0024 |
| LaminDmO | Rabbit |  | 1:200 | Paul Fisher (R836) |
| Puromycin<br>[3RH11] | Mouse | 1:1,000 | 1:50 | Kerafast, #EQ0001 |
| Rps3 | Rabbit | 1:1,000 |  | Bethyl laboratories # A303-840A |

**Supplemental Table 4.**
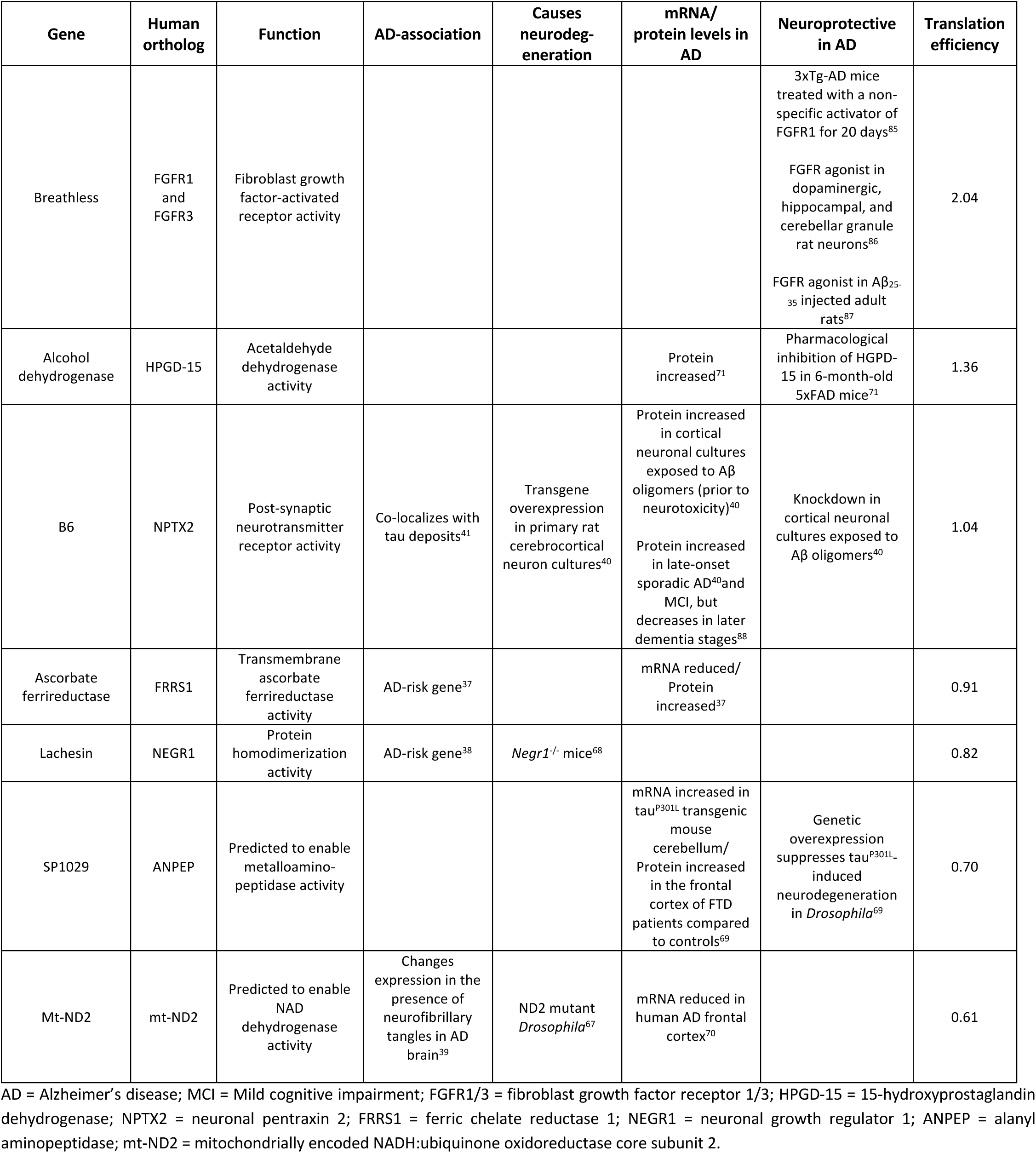
Differentially translated mRNA in tau^R406W^ transgenic *Drosophila* have been implicated in neurodegenerative disorders. Summary of the current literature findings linking the profiling-recognized, efficiently translated genes to Alzheimer’s disease and neurodegeneration.

| Gene | Human ortholog | Function | AD-association | Causes neurodegeneration | mRNA/protein levels in AD | Neuroprotective in AD | Translation efficiency |
| --- | --- | --- | --- | --- | --- | --- | --- |
| Breathless | FGFR1 and FGFR3 | Fibroblast growth factor-activated receptor activity | | | | 3xTg-AD mice treated with a non-specific activator of FGFR1 for 20 days<sup>85</sup><br>FGFR agonist in dopaminergic, hippocampal, and cerebellar granule rat neurons<sup>86</sup><br>FGFR agonist in Aβ<sub>25-35</sub> injected adult rats<sup>87</sup> | 2.04 |
| Alcohol dehydrogenase | HPGD-15 | Acetaldehyde dehydrogenase activity |  |  | Protein increased<sup>71</sup> | Pharmacological inhibition of HPGD-15 in 6-month-old 5xFAD mice<sup>71</sup> | 1.36 |
| B6 | NPTX2 | Post-synaptic neurotransmitter receptor activity | Co-localizes with tau deposits<sup>41</sup> | Transgene overexpression in primary rat cerebrocortical neuron cultures<sup>40</sup> | Protein increased in cortical neuronal cultures exposed to Aβ oligomers (prior to neurotoxicity)<sup>40</sup><br>Protein increased in late-onset sporadic AD<sup>40</sup> and MCI, but decreases in later dementia stages<sup>88</sup> | Knockdown in cortical neuronal cultures exposed to Aβ oligomers<sup>40</sup> | 1.04 |
| Ascorbate ferrioreductase | FRRS1 | Transmembrane ascorbate ferrioreductase activity | AD-risk gene<sup>37</sup> |  | mRNA reduced/ Protein increased<sup>37</sup> |  | 0.91 |
| Lachesin | NEGR1 | Protein homodimerization activity | AD-risk gene<sup>38</sup> | *Negr1*<sup>-/-</sup> mice<sup>68</sup> |  |  | 0.82 |
| SP1029 | ANPEP | Predicted to enable metalloaminopeptidase activity |  |  | mRNA increased in tau<sup>P301L</sup> transgenic mouse cerebellum/ Protein increased in the frontal cortex of FTD patients compared to controls<sup>69</sup> | Genetic overexpression suppresses tau<sup>P301L</sup>-induced neurodegeneration in *Drosophila*<sup>69</sup> | 0.70 |
| Mt-ND2 | mt-ND2 | Predicted to enable NAD dehydrogenase activity | Changes expression in the presence of neurofibrillary tangles in AD brain<sup>39</sup> | ND2 mutant *Drosophila*<sup>67</sup> | mRNA reduced in human AD frontal cortex<sup>70</sup> |  | 0.61 |

AD = Alzheimer's disease; MCI = Mild cognitive impairment; FGFR1/3 = fibroblast growth factor receptor 1/3; HPGD-15 = 15-hydroxyprostaglandin dehydrogenase; NPTX2 = neuronal pentraxin 2; FRRS1 = ferric chelate reductase 1; NEGR1 = neuronal growth regulator 1; ANPEP = alanyl aminopeptidase; mt-ND2 = mitochondrially encoded NADH:ubiquinone oxidoreductase core subunit 2.

## Notes

### Competing Interest Statement

The authors have declared no competing interest.

